# Physical activity phenotyping with activity bigrams, and their association with BMI

**DOI:** 10.1101/121145

**Authors:** Louise AC Millard, Kate Tilling, Debbie A Lawlor, Peter A Flach, Tom R Gaunt

**Affiliations:** MRC Integrative Epidemiology Unit (IEU), University of Bristol, Bristol, United Kingdom; School of Social and Community Medicine, University of Bristol, Bristol, United Kingdom; Intelligent Systems Laboratory, University of Bristol, Bristol, United Kingdom

**Keywords:** Physical activity, bigrams, body mass index, ALSPAC

## Abstract

**Background:** Analysis of physical activity usually focuses on a small number of summary statistics derived from accelerometer recordings: average counts per minute, and the proportion of time spent in moderate-vigorous physical activity or in sedentary behaviour. We show how bigrams, a concept from the field of text mining, can be used to describe how a person’s activity levels change across (brief) time points. These variables can, for instance, differentiate between two people with the same time in moderate activity, where one person often stays in moderate activity from one moment to the next and the other does not.

**Methods:** We use data on 4810 participants of the Avon Longitudinal Study of Parents and Children (ALSPAC). We generate a profile of bigram frequencies for each participant and test the association of each frequency with body mass index (BMI), as an exemplar.

**Results:** We found several associations between changes in bigram frequencies and BMI. For instance, a 1 standard deviation decrease in the number of adjacent minutes in sedentary then moderate activity (or vice versa), with a corresponding increase in the number of adjacent minutes in moderate then vigorous activity (or vice versa), was associated with a 2.36 kg/m^2^ lower BMI [95% CI: -3.47, -1.26], after accounting for the time spent at sedentary, low, moderate and vigorous activity.

**Conclusions:** Activity bigrams are novel variables that capture how a person’s activity changes from one moment to the next. These variables can be used to investigate how sequential activity patterns associate with other traits.

**Key Messages:**

- Epidemiologists typically use only a small number of variables to analyse the association of physical activity with other traits, such as the average counts per minute and the proportion of time spent in moderate-vigorous physical activity or being sedentary.
- We demonstrate how activity bigrams can be used as a set of interpretable variables describing how a person’s activity levels change from one moment to the next.
- Testing the association of activity bigrams with exposures or outcomes can help us gain further understanding of how physical activity is associated with other traits; with further research they might provide evidence for more refined public health advice.

## INTRODUCTION

Physical activity – defined as any bodily movement that results in energy expenditure – is associated with many diseases, such as diabetes (1) and coronary heart disease (2). Research increasingly uses objective measures of physical activity recorded using accelerometers, rather than self-report via questionnaires that are affected by reporting bias and measurement error (3). Cohort participants wear an accelerometer that measures accelerations at time intervals typically ranging from 0.01 seconds (4) to 1 minute (5,6). This high-resolution time-series data potentially contains much valuable information about a person’s activity. To date, however, only a small number of variables derived from accelerometer recordings have been used: average counts per minute (mCPM), and the proportion of time spent in moderate-vigorous physical activity (MVPA) or sedentary behaviour (SB) (5,7–12). These measures only include a fraction of the information contained in accelerometer sequences, and this may lead to bias and a loss of power when using these variables as measures of physical activity. As public health advice has, to date, been informed by research using these limited variables, it is possible that analyses using other aspects of this data would support more refined advice.

Recently, work has been conducted to generate other variables describing physical activity. Goldsmith et al. used functional data analyses to model diurnal physical activity profiles, and test the association of these profiles with other traits (13). For example, they identified that during daytime hours girls are less active than boys, but this difference is not present in the evening. Evenson et al. used latent class analysis to assign participants to groups based on their activity levels across a one week period, to identify common weekly patterns of activity (14). Their analyses using MVPA identified an interesting set of 4 groups – two lower activity groups with stable MVPA across the week, and two higher activity groups, one most active between Monday and Thursday and the other most active between Friday and Sunday. Augustin et al. used a histogram of activity counts as a functional summary of activity (15), which is beneficial because it does not assume that the association of activity on a second trait across activity levels is linear, and also allows this assumption to be tested. Their analysis with fat mass found a non-linear association across activity intensities.

One key aspect of accelerometer sequences not captured by mCPM and time spent in different intensities of activity from sedentary to MVPA, or the more recent methods described above, is variability in a person’s activity levels from one moment to the next. For instance, two people may have the same mCPM and also the same total time spent in MVPA, but the first person may stay at the vigorous activity level for one continuous period, whereas the second enters into the vigorous activity state for more frequent, shorter bouts. It is increasingly recognised that variability of a trait about the mean level can have important associations with exposures and outcomes, independently of the mean level (e.g. variation of systolic blood pressure (16–18)).

At present, most physical activity guidelines recommend accumulating at least 150 minutes of moderate intensity activity or 75 minutes of vigorous intensity activity a week (19,20), with no advice on possible benefits of time-varying intensities. There is increasing interest in the possible health benefits of undertaking repeat short bursts of high intensity activity, referred to as high intensity interval training (HIIT) (21–23). While HIIT research assesses the benefits of short periods of very high intensity activity, there are many other sequential activity patterns that might also be beneficial (or detrimental) to a person’s health. However, methods for assessing the association of a sequence of exposures (rather than the mean level) with an outcome are not widely used.

Sequential data, like that from accelerometers, occurs in many settings. The field of text mining seeks to learn models to make predictions from the sequence of words in a document (24). A common approach is to treat each document as an unordered collection of words, each called a unigram, and to use the frequency of each word in the document as a variable in analyses. The set of words and associated frequencies is known as a bag of words. This is equivalent to the way epidemiologists treat accelerometer data, after the sequence is first categorised into sedentary, low and moderate/vigorous activity. The accelerometer sequence is treated as an unordered collection of these activity categories and the variables MVPA and SB denote the frequency (or proportion) of each category in the sequence. Hence we can view these activity categories as *activity unigrams*, and the set of activity unigrams with the associated frequencies as a bag of activity unigrams. While a unigram is a sequence of length one, this can be generalised to *n*-grams – sequences of length *n* – and bags of n-grams (see Supplementary material section S1 for examples). This provides opportunities to extend representations of physical activity beyond activity unigrams.

In this paper we use 2-grams, referred to as bigrams, to represent sequential patterns in a person’s accelerometer sequence. This is useful as we can then ask, for instance, how often is a person in the moderate state at time *t* and the vigorous state at time *t+1*? Activity bigrams can be used to examine how changes in activity from one moment to the next associates with other traits. We demonstrate our novel approach with BMI, as an exemplar.

## METHODS

### Participants

We used data on participants in the Avon Longitudinal Study of Parents and Children (ALSPAC), a prospective population-based cohort. The ALSPAC study website contains details of all the data that are available through a fully searchable data dictionary: [<u>http://www.bris.ac.uk/alspac/researchers/data-access/data-dictionary/</u>]. The study methods are described in detail elsewhere (25). In brief, ALSPAC recruited 14,541 pregnant women resident in Avon, United Kingdom, with expected dates of delivery between 1 April 1991 and 31 December 1992 (<u>http://www.alspac.bris.ac.uk</u>). These mothers and their children have been followed with regular assessments since this time. Ethical approval for the study was obtained from the ALSPAC Ethics and Law Committee and the Local Research Ethics Committees.

### Data collection

Physical activity was measured using the uni-axial Actigraph 7164 accelerometer that measures vertical accelerations. All children who attended the age 11 clinic were asked to wear an accelerometer on their waist for 7 days, taking it off while sleeping, showering, bathing or swimming. The devices were programmed to start recording at 05:00 a.m. the day after the clinic. The sum of activity counts (a measure of acceleration) over each one-minute epoch (interval) was recorded, giving a maximum sequence of 10 080 values for each participant. We refer to each two-minute interval in a person’s sequence as an epoch pair. The total number of epoch pairs in a sequence is equal to the length of the sequence minus one.

Weight and height were measured at the age 11 clinic, with the child in light clothing without shoes. BMI was calculated as weight in kilograms divided by height in meters squared. We consider the following potential confounders: child gender, exact age at age 11 clinic, parity, household social class, maternal education, maternal smoking during pregnancy and child ethnicity (details of how these were assessed are provided in Supplementary material section S2).

### Study sample

Of the 6080 participants with accelerometer data we excluded from analyses 68 participants who did not have 7 days of recorded accelerometer data. We further excluded 40 participants with no measure of BMI. We assumed that continuous sequences of zero activity counts of length greater than 60 epochs (1 hour) meant that the participant was not wearing their device and treated such periods as missing accelerometer data (26). Accelerometer data was considered invalid if: 1) there were fewer than three valid days, where a valid day is defined as at least 8 hours of wear time, or 2) the average activity level per minute was greater than 1500, as this was deemed infeasible. We excluded 240 participants with invalid accelerometer data. We removed 66 participants who were siblings to other participants in this sample. We excluded a further 856 participants with no value for at least one confounding factor, giving a resultant sample size of 4810 participants. A participant flow diagram is given in Figure 1.

**Figure 1:**
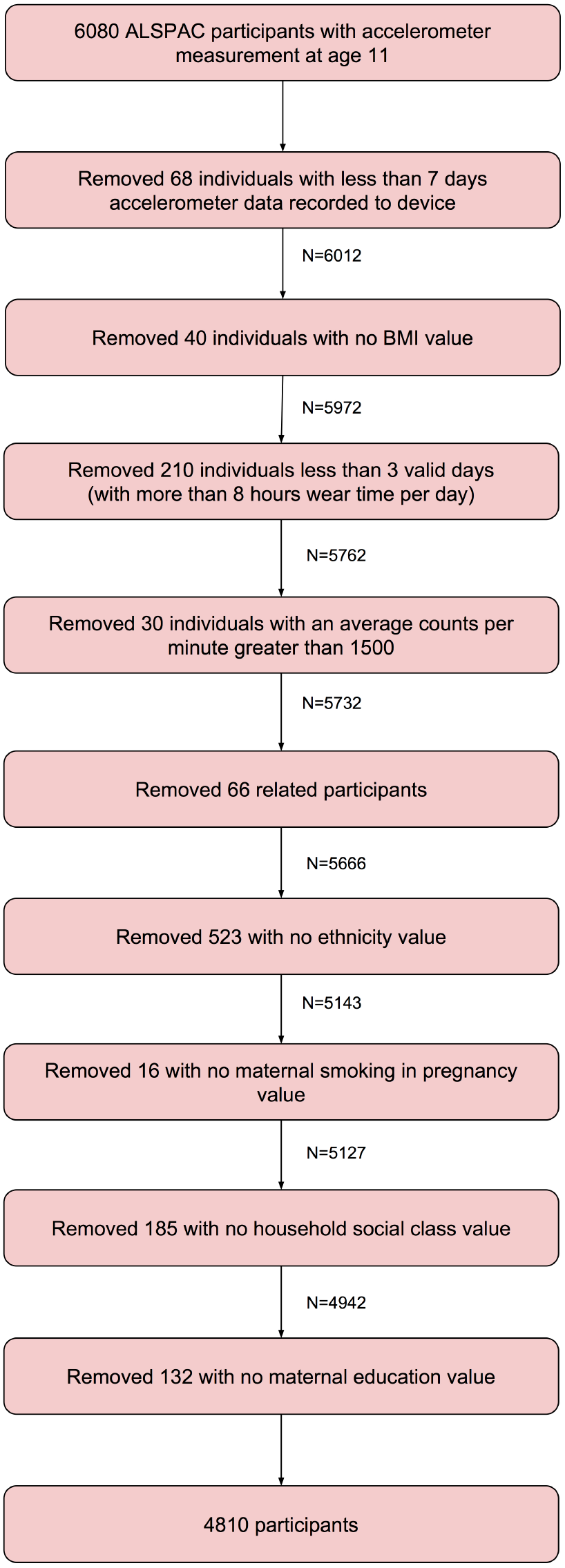
**Participant flow diagram showing creation of our sample in ALSPAC**

### Statistical analyses

The activity levels at each epoch of participant’s accelerometer sequences (excluding non-wear time) were categorized into four groups of activity intensities: sedentary: 0-100 activity counts per minute; low: 101-2019; moderate: 2020-5998, and vigorous: 5999+ (6,26), denoted S, L, M and V, respectively. We refer to these as activity *states*, to distinguish from the continuous activity *levels* of the original accelerometer data.

#### Relating mean activity levels to outcomes

We use univariate linear regression (`regress` function in Matlab) to test the association of the mean activity levels per minute over the time where a participant wore the accelerometer (mCPM) and the variance of these activity levels per minute (vCPM), individually, with BMI. We test this association before and after adjustment for potential confounding factors, and also after mutual adjustment of mCPM and vCPM in order to examine their independent associations with BMI.

Within one day there are a finite number of occurrences of activity states in total (the number of minutes = 1440), such that as the frequency of one activity state increases, this must be coupled with a decrease in frequency of at least one other activity state. This means that an increase in frequency of the moderate state may, for instance, be associated with lower BMI when the additional frequency comes from the sedentary state, but not when it comes from the vigorous state. We calculate the average number of minutes each participant spends in S, L, M and V activity states per day, denoted *S_d_, L_d_, M_d_* and *V_d_*, respectively. We then use univariate linear regression to estimate the association of transferring time between pairs of activity states, with BMI. We assign, in turn, one activity state as a baseline and another as a comparison, and calculate the total remaining time per day. The comparison state and remaining time are included in the model and the baseline is not included. For example, we use the following model and multiply *β*_1_ by 10 to estimate the difference in means of BMI for a 10-minute per day transfer from the sedentary (baseline) to the moderate (comparison) activity state:

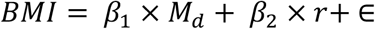

where *r* = *n_d_* − (*M_d_* + *S_d_*) and *n_d_* is the number of epochs per day (in this case 1440). We use the number of minutes spent in each activity state in our models rather than the proportion of non-missing time, as we are interested in how the actual amount of time spent in each state associates with BMI. We note that swapping the baseline and comparison activity states results in a reciprocal model with estimate –*β*_1_. We test these associations both before and after adjustment for potential confounders.

#### Modelling activity sequences with activity bigrams

We derive a set of variables denoting the number of times a particular bigram occurs in a person’s sequence, on average per day. Given the four activity states – sedentary, low, moderate and vigorous, there are 16 bigrams: SS, SL, SM, SV, LS, LL, LM, LV, MS, ML, MM, MV, VS, VL, VM and VV. For instance, SL denotes the occurrence of the sedentary state at time *t*, followed by the low state at time *t+1*. In this work we use a one-minute epoch such that *t*=1 minute. Formally, the frequency of a bigram AB per day is given by:

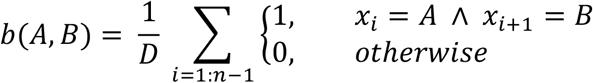

for number of days *D* (in this work *D=7*) and sequence x_i_ ={x_1_, x_2_ … x_n_} where *x ε {S,L,M,V}*. Figure 2 provides two example sequences where the values of the common activity statistics (MVPA, SB and mCPM) are the same, but the frequency of bigrams differ. For example, the SS bigram occurs three times in sequence A but only once in sequence B. Epoch pairs are overlapping such that a bigram occurring in the time period [t, t+1] overlaps with the bigrams at [t-1, t] and [t+1, t+2]. This means that the frequency of each bigram in a sequence does not correspond to a specific amount of time. For instance, the sequences SSSLS and SSLSS both have two occurrences of the SS bigram, but over three and four minutes, respectively.

**Figure 2:**
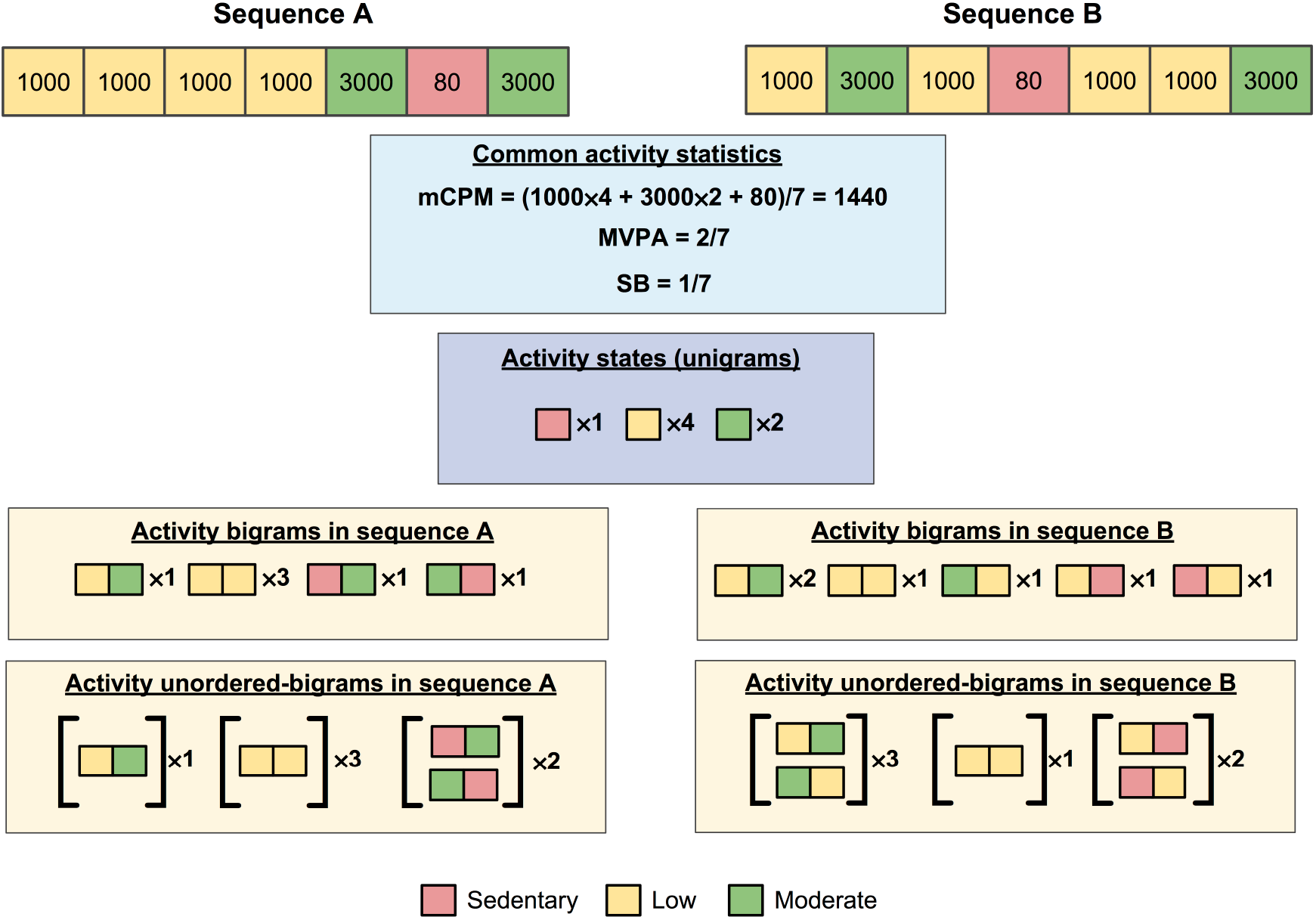
Illustrative examples of common physical activity statistics and our novel activity bigrams. mCPM: average counts per minute; MVPA: proportion of time spent in moderate-vigorous physical activity; SB: sedentary behaviour. Illustration shows two 7-minute activity sequences, where each coloured block denotes a one-minute interval with a given activity level. Sequence A and sequence B have the same number of occurrences of each activity state (with the same activity levels) and so have the same values for the common activity statistics and frequency of each activity state, but the different order of activity states means they have different frequencies of bigrams and unordered-bigrams.

#### Relating activity bigrams to outcomes

As with activity states, a person can only have a fixed number of occurrences of bigrams in total per day, such that as the frequency of one bigram increases the frequency of at least one other must decrease. Also, because bigrams are overlapping, a change of an epoch pair in a sequence from one bigram to another will often change the number of occurrences of at least one other bigram, and these changes depend on the particular sequence (see examples in Figure 3 and supplementary section S3). For these reasons, we investigate how BMI changes as the average frequency of bigrams per day increases for one bigram while at the same time decreasing for another bigram, while allowing for collateral changes in other bigrams.

**Figure 3:**
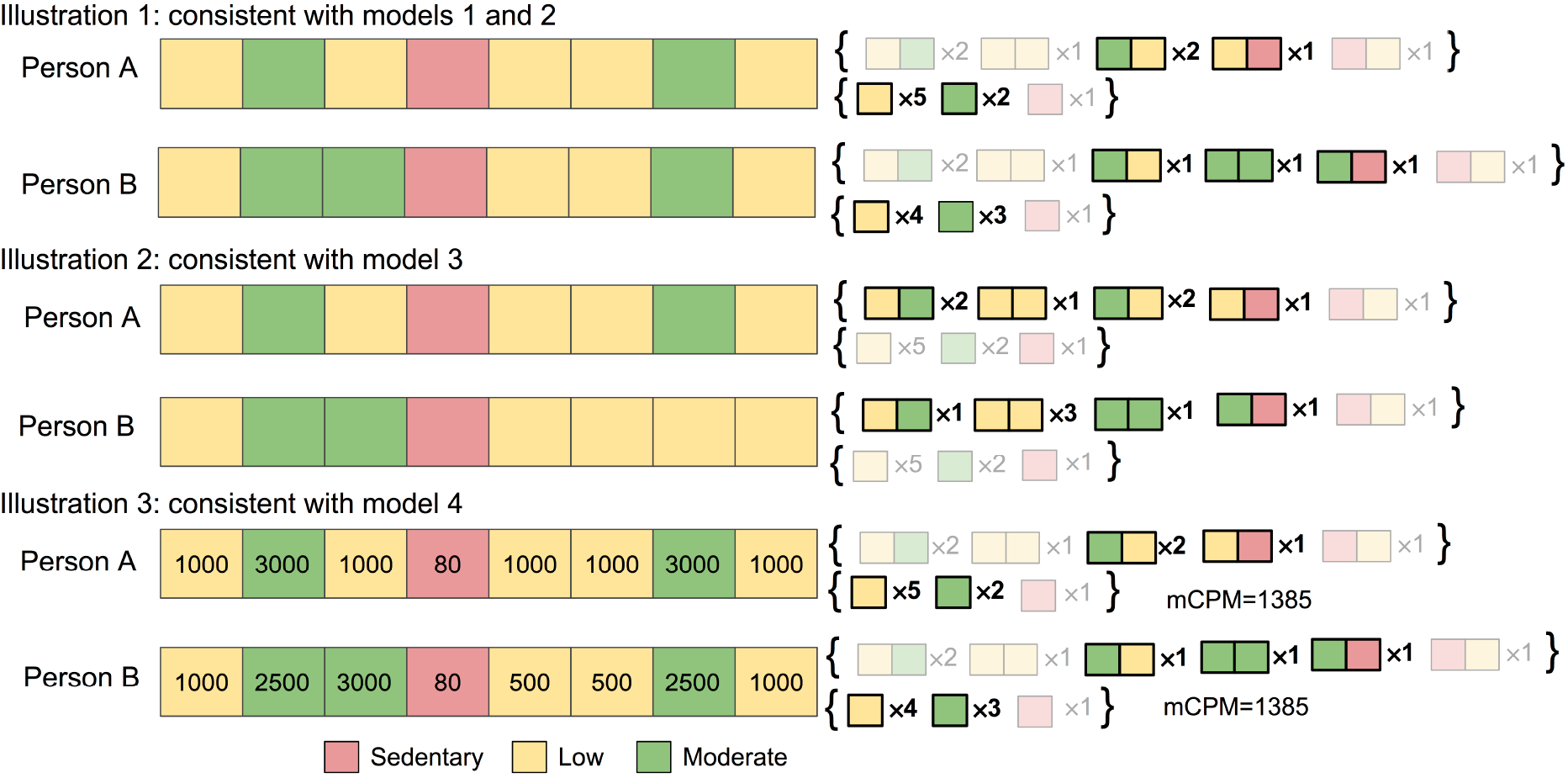
Illustrative examples of real differences in activity sequences consistent with our models, for baseline ML (moderate followed by low) and comparison MM (moderate followed by moderate) activity bigrams. mCPM: average counts per minute. Each illustration shows the activity of two people during an 8-minute period, where each coloured block denotes a one-minute interval. Illustration 3 also shows the activity level of each minute. Curly brackets on right hand side show the frequency of each activity bigram and activity state, where those emboldened have different values for person A and B. Illustration 1: Consistent with models 1 and 2 because swapping ML (the moderate followed by low bigram) with MM (the moderate followed by moderate bigram) increases the occurrence of MM and decreases the occurrence of ML by the same amount. Illustration 2: Consistent with model 3 because: 1) swapping ML with MM increases the occurrence of MM and decreases the occurrence of ML by the same amount, and 2) the time spent in sedentary, low, moderate and vigorous does not change. Illustration 3: Consistent with model 4 because: 1) swapping ML with MM increases the occurrence of MM and decreases the occurrence of ML by the same amount, and 2) the average counts per minute (mCPM) does not change.

We use univariate linear regression and assign one bigram as a baseline (i.e. not included in the model) and another as a comparison (i.e. included in the model), and adjust for the remaining number of bigrams in a day. For example, we use the following model and multiply ***β*_1_** by 10 to estimate the difference in means of BMI for a 10 epoch pair increase of the SL bigram, coupled with a 10 epoch pair decrease of the SS bigram:

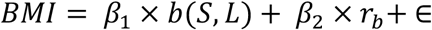

where the remaining number of bigrams in a day is given by: ***r_b_* = *n_d_* − 1 − *b*(*S, L*) − *b*(*S, S*)**. We work with 10 epoch pairs as a non-negligible amount for which a person may reasonably be expected to change their activity. Adjusting for the combined frequency of the remaining bigrams rather than the frequency of each bigram separately, allows for collateral changes in the frequency of these bigrams (while their total frequency remains the same). While we may think of these models as representing a swap from one bigram to another at particular positions in a person’s sequence, in fact any increase in frequency of one bigram that is accompanied by an *equal* decrease in frequency of another bigram, is consistent with these models (see Supplementary section S3 for further explanation).

We investigated the impact of potential confounding by characteristics that relate to both the bigram measures and BMI by including these characteristics as covariables in the regression analyses (see **Table 1** for confounders). We also considered that the following accelerometer variables might confound associations of bigrams with BMI: mCPM and the number of minutes spent in sedentary, low, moderate or vigorous activity. This is because these will be related to the bigrams and it is well established that they are related to BMI. These adjustments were made in a series. In all analyses we undertook the following:

**Table 1:** Summary statistics of ALSPAC participants who attended the focus@11 clinic, who are included and not included in our sample

|  | Attended focus at 11 clinic and not in sample |  | Attended focus at 11 clinic and in sample |  | Difference between participants and non-participants<sup>2</sup> |
| --- | --- | --- | --- | --- | --- |
|  | Number of participants | Mean (SD) or N (%)<sup>1</sup> | Number of participants | Mean (SD) or N (%)<sup>1</sup> | Odds ratio [95% CI] |
| BMI in kg/m<sup>2</sup> (mean (SD)) | 2297 | 19.39 (3.73) | 4810 | 18.97 (3.30) | 0.97 [0.95, 0.98] |
| **Potential confounding factors** |  |  |  |  |  |
| Age in years at age 11 clinic (mean (SD)) | 2343 | 11.82 (0.26) | 4810 | 11.77 (0.23) | 0.45 [0.36, 0.55] |
| Parity (%) - 0 | 2343 | 869 (37.09) | 4810 | 2219 (46.13) | 0.66 [0.62, 0.70] |
| 1 |  | 619 (26.42) |  | 1688 (35.09) |  |
| 2+ |  | 855 (36.49) |  | 903 (18.77) |  |
| Sex - % female | 2343 | 1114 (47.55) | 4810 | 2552 (52.43) | 1.22 [1.10, 1.34] |
| Ethnicity - % non-white | 1640 | 82 (5.00) | 4810 | 169 (3.51) | 0.69 [0.53, 0.91] |
| Mother smokes in pregnancy - % yes | 1900 | 455 (23.95) | 4810 | 821 (17.07) | 0.65 [0.57, 0.74] |
| Household social class (%) – I | 1440 | 199 (13.82) | 4810 | 776 (16.13) | 0.86 [0.81, 0.91] |
| II |  | 614 (42.64) |  | 2185 (45.43) |  |
| III (non-manual) |  | 358 (24.86) |  | 1215 (25.26) |  |
| III (manual) |  | 180 (12.50) |  | 468 (9.73) |  |
| IV/V |  | 89 (6.18) |  | 166 (3.45) |  |
| Maternal education (%) - Less than O level | 1503 | 322 (21.42) | 4810 | 877 (18.23) | 1.06 [1.00, 1.13] |
| % O level |  | 530 (35.26) |  | 1788 (37.17) |  |
| % A level |  | 418 (27.81) |  | 1333 (27.71) |  |
| % Degree |  | 233 (15.50) |  | 812 (16.88) |  |

BMI: body mass index; SD: standard deviation.
<sup>1</sup> Mean (SD) for continuous and percentage for binary variables.
<sup>2</sup> Odds ratio for participants included in our sample versus participants who attended the age 11 clinic but are not in our sample (reference group), for a 1 unit increase in continuous variable (using variable units as described in column 1), or comparison group (indicated in column 1) versus baseline group for binary variables, or a 1 category increase for ordinal categorical variables. For example, an odds ratio of 0.97 [0.95, 0.98] for BMI means that on average a participant (who attended the age 11 clinic) is 3% [95% CI: 5%, 2%] less likely to be in our sample for each 1kg/m<sup>2</sup> increase in BMI.

- model 1 – unadjusted;
- model 2 – adjusted for child gender, exact age at age 11 clinic, parity, household social class, maternal education, maternal smoking during pregnancy and child ethnicity;
- model 3 – as model 2 and additionally adjusted for the mean number of minutes per day spent in the activity states: sedentary, low, moderate and vigorous;
- model 4 – as model two and additionally adjusted for mCPM.

Figure 3 provides illustrative examples showing how our models relate to differences in activity sequences (see Supplementary material section S3 and Supplementary table 1 for further examples).

#### Relating unordered-bigrams to outcomes

While bigrams denote the ordered occurrence of consecutive activity states it may be the case that it is the adjacent occurrence of activity states that matters rather than the sequential order. For instance, the frequency of the MV bigram may associate with BMI because M and V are adjacent rather than because V follows M, and where this is true we would expect the associations for MV with BMI to be comparable to the associations for VM with BMI.

We repeat our analyses using an unordered version of bigrams, in order to maximise the power of our analyses, and refer to these as unordered-bigrams (u-bigrams). Given the activity states (sedentary, low, moderate and vigorous), there are 10 u-bigrams: [SS], [SL], [SM], [SV], [LL], [LM], [LV], [MM], [MV] and [VV]. For instance, the [SL] u-bigram corresponds to the bigrams SL and LS, and [VV] corresponds to the bigram VV. Formally the frequency of a u-bigram per day is calculated as:

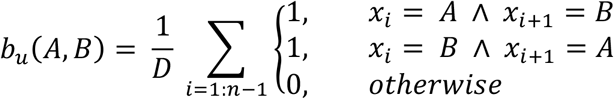

for u-bigram [AB], number of days *D*, and sequence x_i_ ={x_1_, x_2_ … x_n_} where *x ε {S,L,M,V}*. Clearly, the u-bigrams [SS], [LL], [MM], and [VV] are equivalent to the bigrams SS, LL, MM, and VV, respectively. We refer to two bigrams AB and BA, corresponding to the u-bigram [AB], as the reciprocal of each other. Example u-bigrams are given in Figure 2 and example sequence changes consistent with our models are given in Supplementary table 2.

Summary statistics (Table 1) were generated using Stata SE14. All other analyses are performed in Matlab (R2015). All code is available at [https://github.com/MRCIEU/activityBigrams/]. Git tag v0.1 corresponds to the version presented here.

## RESULTS

Table 1 shows characteristics of participants included in our analysis sample compared with those who were eligible (i.e. attended the age 11 clinic) but were not included in our sample because of missing accelerometer, BMI or confounder data. Participants who were younger, lighter, female, white, with a higher household social class (nearer to class I), higher maternal education and whose mothers did not smoke in pregnancy were more likely to be in our sample (than have attended the same clinic but not be in our sample), though the magnitudes of these differences were small.

### Associations of conventional summary activity variables (mCPM and time spent in activity states), with BMI

Before presenting the results of our novel activity bigram variables in the following section, here we present results for the common activity statistics (mCPM and time spent in activity states: sedentary, low, moderate and vigorous) and vCPM. Table 2 and Figure 4 show the association of mCPM and vCPM with BMI. The variables mCPM and vCPM are strongly correlated (Pearson correlation coefficient =0.69). After adjustment for confounders a 100 count per minute increase in mCPM is associated with a 0.283 kg/m^2^ lower BMI [95% CI: - 0.337, -0.229], and after adjustment for vCPM this association remains. After adjustment for confounders a 1 SD increase in vCPM is associated with a 0.356 kg/m^2^ lower BMI [95% CI: - 0.449, -0.263]. However, after adjustment for mCPM this association attenuates to the null.

**Figure 4:**
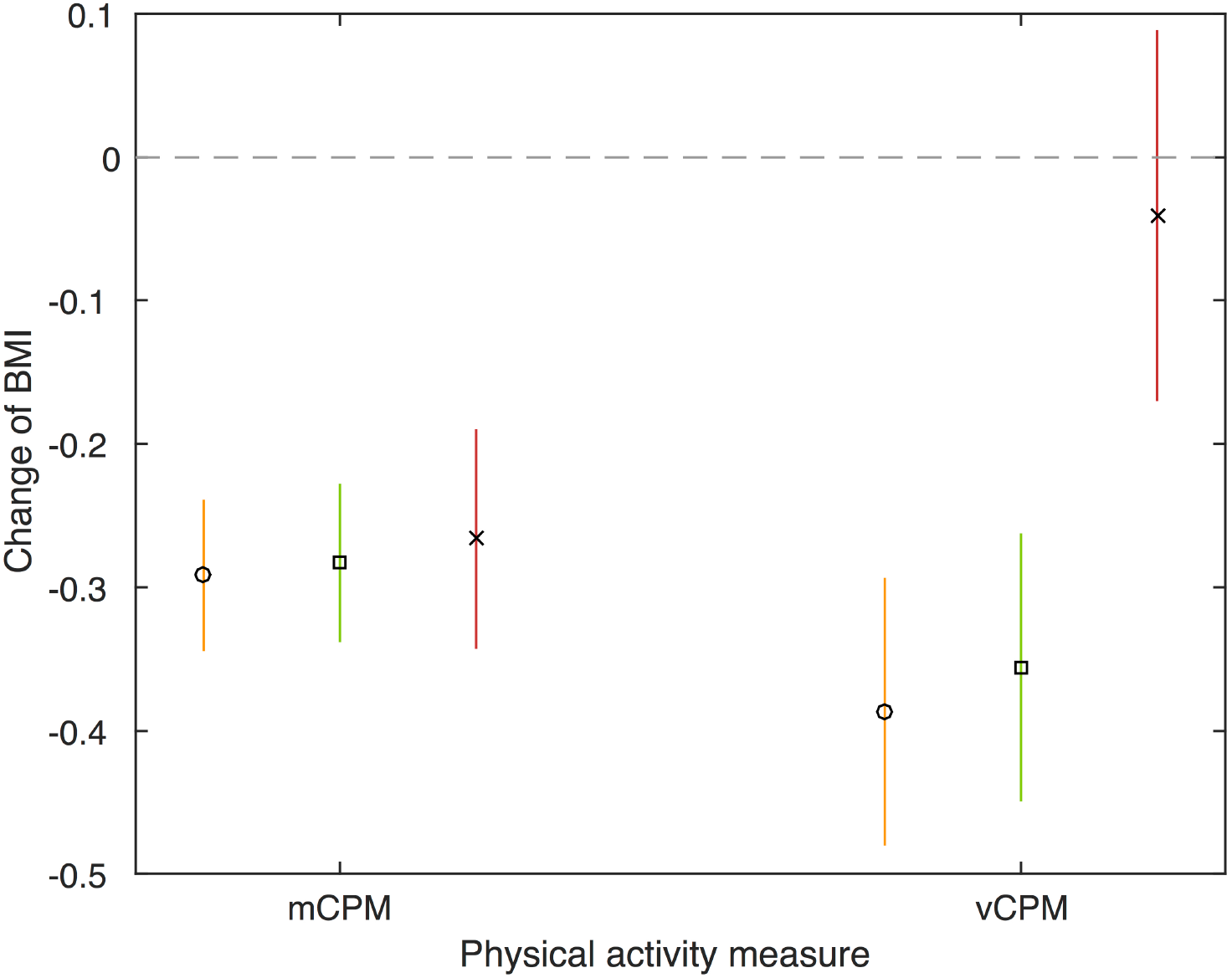
Association of the average counts per minutes and variance of counts per minute with BMI. BMI: body mass index; mCPM: average counts per minute; vCPM: variance of counts per minute. Model 1 (orange circle): unadjusted. Model 2 (green square): adjusted for potential confounders (gender, exact age at age 11 clinic, parity, household social class, maternal education, maternal smoking during pregnancy and child ethnicity). Model 3 (red cross): adjusted for potential confounders (gender, exact age at age 11 clinic, parity, household social class, maternal education, maternal smoking during pregnancy and child ethnicity), and mutually adjusted (vCPM is adjusted for mCPM and vice versa). N=4810.

**Table 2:** **Association of the average counts per minutes and variance of counts per minute with BMI**

| | Difference in means of BMI ($\text{kg/m}^2$) per unit increase in each activity variable [95% confidence interval] N=4810 | | |
| --- | --- | --- | --- |
|  | Model 1 (unadjusted) | Model 2 (non-accelerometer confounder adjusted) | Model 3 (non-accelerometer confounder adjusted and mutually adjusted) |
| mCPM (per 100 counts) | -0.292 [-0.344, -0.240] | -0.283 [-0.337, -0.229] | -0.266 [-0.342, -0.191] |
| vCPM (per 1 SD; $1\text{SD} = 1.06 \times 10^6 \text{ counts}^2$) | -0.387 [-0.480, -0.294] | -0.356 [-0.449, -0.263] | -0.041 [-0.170, 0.088] |

BMI: body mass index; mCPM: average counts per minute; vCPM: variance of counts per minute.
Model 1: unadjusted.
Model 3: adjusted for potential confounders (gender, exact age at age 11 clinic, parity, household social class, maternal education, maternal smoking during pregnancy and child ethnicity), and mutually adjusted (vCPM is adjusted for mCPM and vice versa).

Table 3 and Figure 5 show the unadjusted and confounder adjusted associations of transferring time between activity states (sedentary, low, moderate and vigorous), with BMI. In general transferring time to a higher activity state was associated with a lower BMI, and the greater the increase in activity state, the greater the change in BMI. For example, transferring 10 minutes of time from the sedentary to the vigorous activity state per day was associated with a 0.960 kg/m^2^ lower BMI [95% CI: -1.169, -0.751], after adjustment for non-accelerometer confounders (i.e. Model 2).

**Figure 5:**
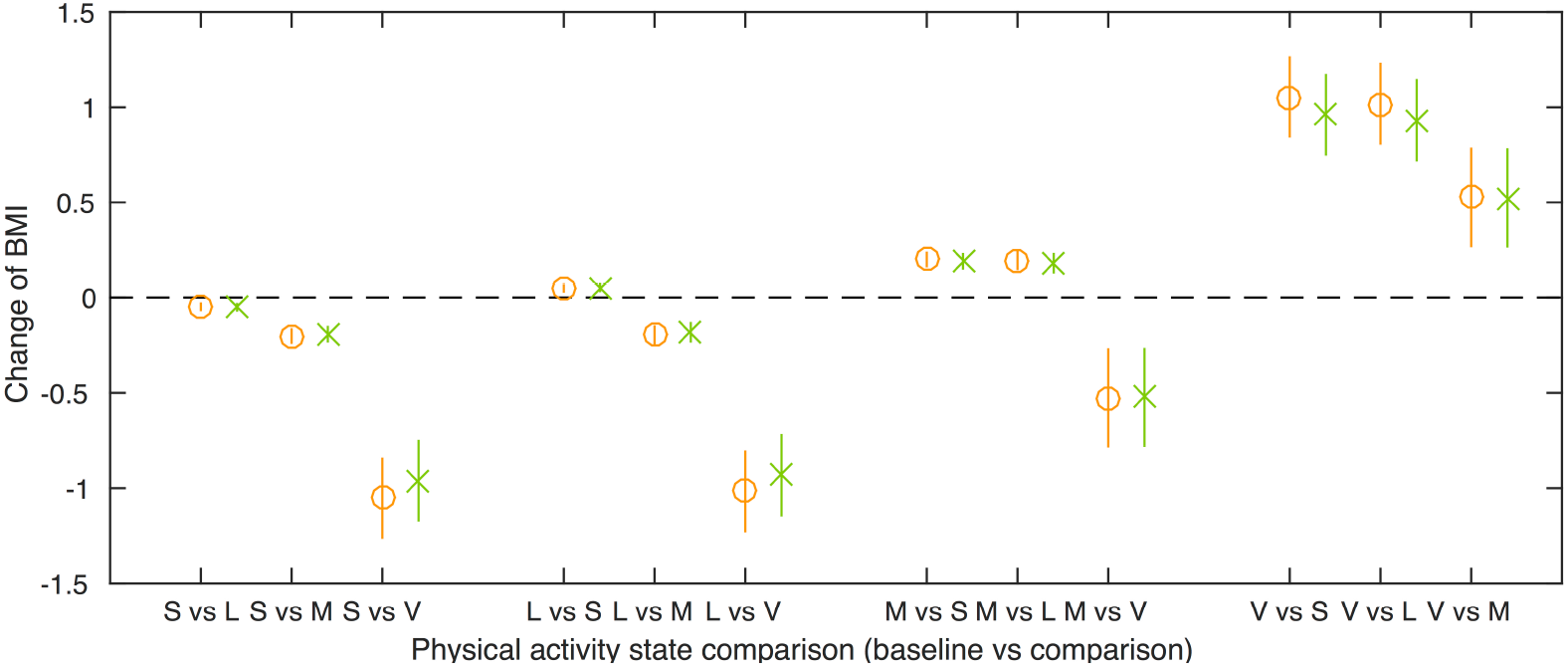
Difference in means of BMI for a 10 occurrence per day transfer from baseline activity state to comparison activity state. BMI: body mass index; S: sedentary; L: low; M: moderate; V: vigorous. Model 1 (orange circle): unadjusted. Model 2 (green cross): adjusted for potential confounders (gender, exact age at age 11 clinic, parity, household social class, maternal education, maternal smoking during pregnancy and child ethnicity). N=4810

**Table 3:**
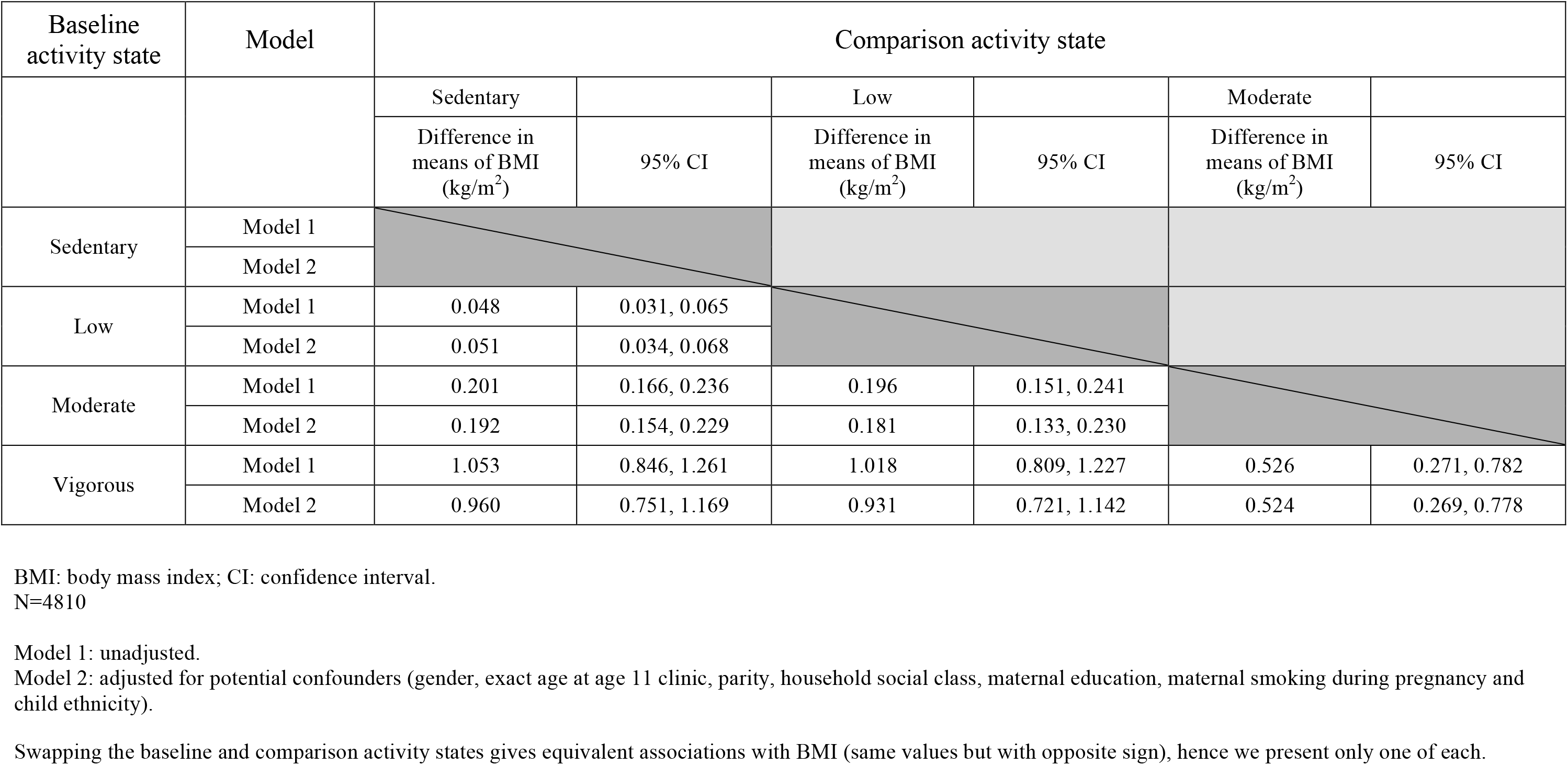
**Difference in means of BMI for 10-minute per day transfer from baseline activity state to comparison activity state**

| Baseline activity state | Model | Comparison activity state |  |  |  |  |  |
| --- | --- | --- | --- | --- | --- | --- | --- |
|  |  | Sedentary |  | Low |  | Moderate |  |
|  |  | Difference in means of BMI (kg/m<sup>2</sup>) | 95% CI | Difference in means of BMI (kg/m<sup>2</sup>) | 95% CI | Difference in means of BMI (kg/m<sup>2</sup>) | 95% CI |
| Sedentary | Model 1 |  |  |  |  |  |  |
|  | Model 2 |  |  |  |  |  |  |
| Low | Model 1 | 0.048 | 0.031, 0.065 |  |  |  |  |
|  | Model 2 | 0.051 | 0.034, 0.068 |  |  |  |  |
| Moderate | Model 1 | 0.201 | 0.166, 0.236 | 0.196 | 0.151, 0.241 |  |  |
|  | Model 2 | 0.192 | 0.154, 0.229 | 0.181 | 0.133, 0.230 |  |  |
| Vigorous | Model 1 | 1.053 | 0.846, 1.261 | 1.018 | 0.809, 1.227 | 0.526 | 0.271, 0.782 |
|  | Model 2 | 0.960 | 0.751, 1.169 | 0.931 | 0.721, 1.142 | 0.524 | 0.269, 0.778 |

BMI: body mass index; CI: confidence interval. N=4810
Model 1: unadjusted.
Model 2: adjusted for potential confounders (gender, exact age at age 11 clinic, parity, household social class, maternal education, maternal smoking during pregnancy and child ethnicity).
Swapping the baseline and comparison activity states gives equivalent associations with BMI (same values but with opposite sign), hence we present only one of each.

### Associations between sequences of physical activity (bigrams and u-bigrams), with BMI

The distributions of the bigrams are shown in Supplementary figure 1. The frequencies of reciprocal bigrams in an individual’s sequence were highly correlated (Pearson correlation coefficients range from 0.479 [95% CI: 0.457, 0.500] for SV versus VS, to 0.999 [95% CI: 0.999, 0.999] for SL versus LS); see Supplementary table 3). The associations of reciprocal bigrams (such as MV and VM) with BMI were largely consistent (see Supplementary table 4 and Supplementary figures 2-17 for bigram results). For example, a 10 epoch pair higher frequency of MV, coupled with a 10 epoch pair lower frequency of SS is associated with a 2.308 kg/m^2^ lower BMI [95% CI: -3.552, -1.064], and a 10 epoch pair higher frequency of VM, coupled with a 10 epoch pair lower frequency of SS is associated with a 1.926 kg/m^2^ lower BMI [95% CI: -3.169, -0.683], after adjustment for confounders and the average frequency of activity states per day (model 3). This suggests that the order of the activity states within a bigram (eg. MV versus VM) does not affect its association with BMI, and so we present the u-bigram results as the main results.

Table 4 and Figure 6 show the associations of frequency changes of u-bigrams, with BMI (models 1, 2 and 4 are shown in Supplementary table 5). An increase in frequency of the [MV] u-bigram, when coupled with a decrease in frequency of all other u-bigrams except [VV], show negative associations with BMI, after adjusting for confounders and both the time spent in each activity state, and mCPM, respectively. For example, a 10 epoch pair higher frequency of [MV], coupled with a 10 epoch pair lower frequency of [SM] is associated with a 1.840 kg/m^2^ lower BMI [95% CI: -2.701, -0.980], after adjustment for confounders and the time spent in each activity state. We also found associations for an increase in frequency of the [MM] u-bigram, when coupled with a decrease in frequency of the [SS] and [LL] u-bigrams, that remained after adjusting for the both the time spent in each activity state, and mCPM, respectively.

**Figure 6:**
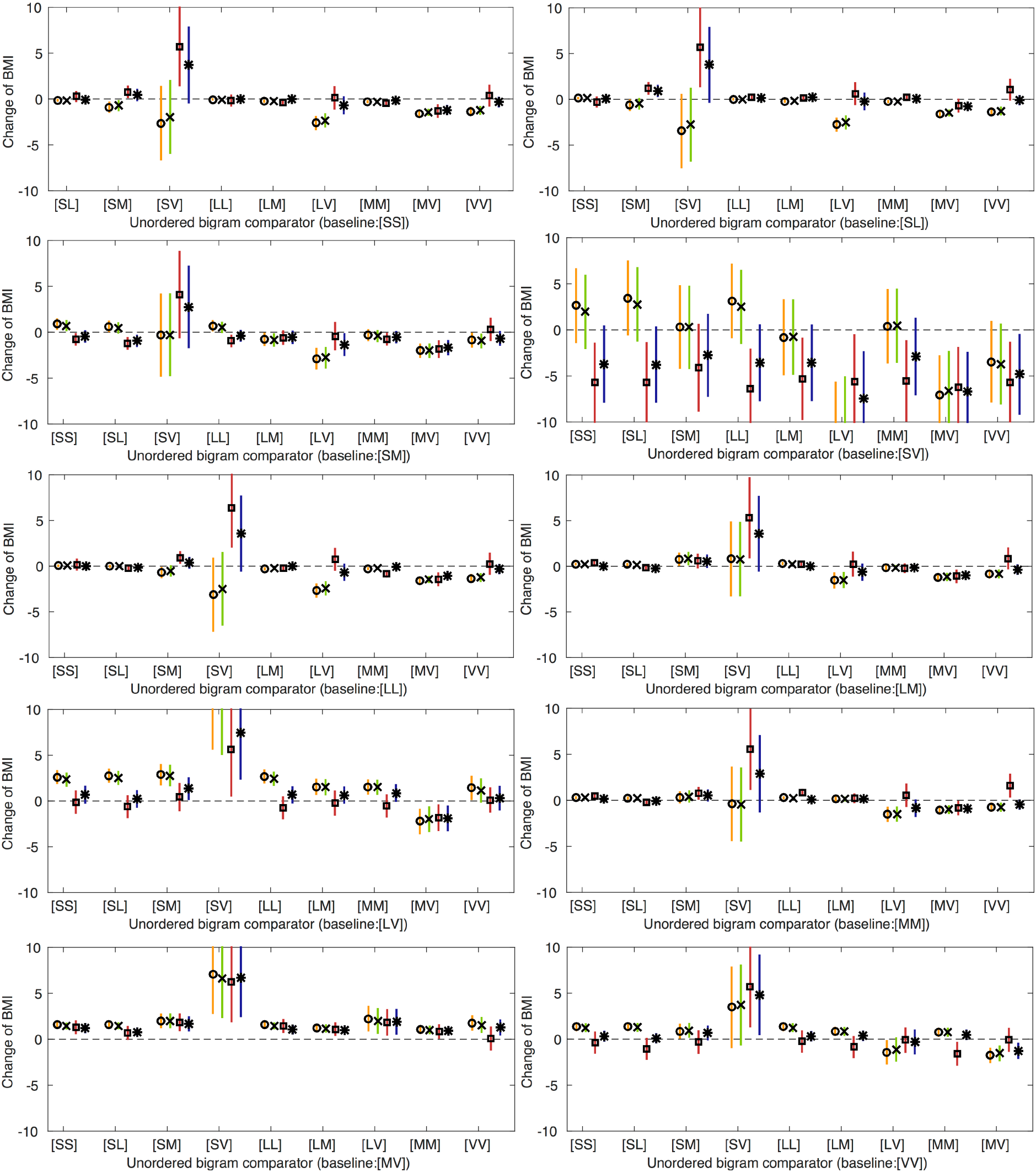
Difference in means of BMI for a 10 epoch pair lower frequency of baseline activity u-bigram, coupled with a 10 epoch pair higher frequency of comparison activity u-bigram. BMI: body mass index; u-bigram: unordered-bigram; S: sedentary; L: low; M: moderate; V: vigorous. Swapping the comparison and baseline u-bigram gives equivalent estimates of association with BMI (same values but with opposite sign). Model 1 (circle): unadjusted. Model 2 (cross): adjusted for potential confounders (gender, exact age at age 11 clinic, parity, household social class, maternal education, maternal smoking during pregnancy and child ethnicity). Model 3 (square): adjusted for potential confounders (gender, exact age at age 11 clinic, parity, household social class, maternal education, maternal smoking during pregnancy and child ethnicity) and activity states (time spent in sedentary, low, moderate and vigorous activity). Model 4 (star): adjusted for potential confounders (gender, exact age at age 11 clinic, parity, household social class, maternal education, maternal smoking during pregnancy and child ethnicity), and mCPM. N=4810

**Table 4:**
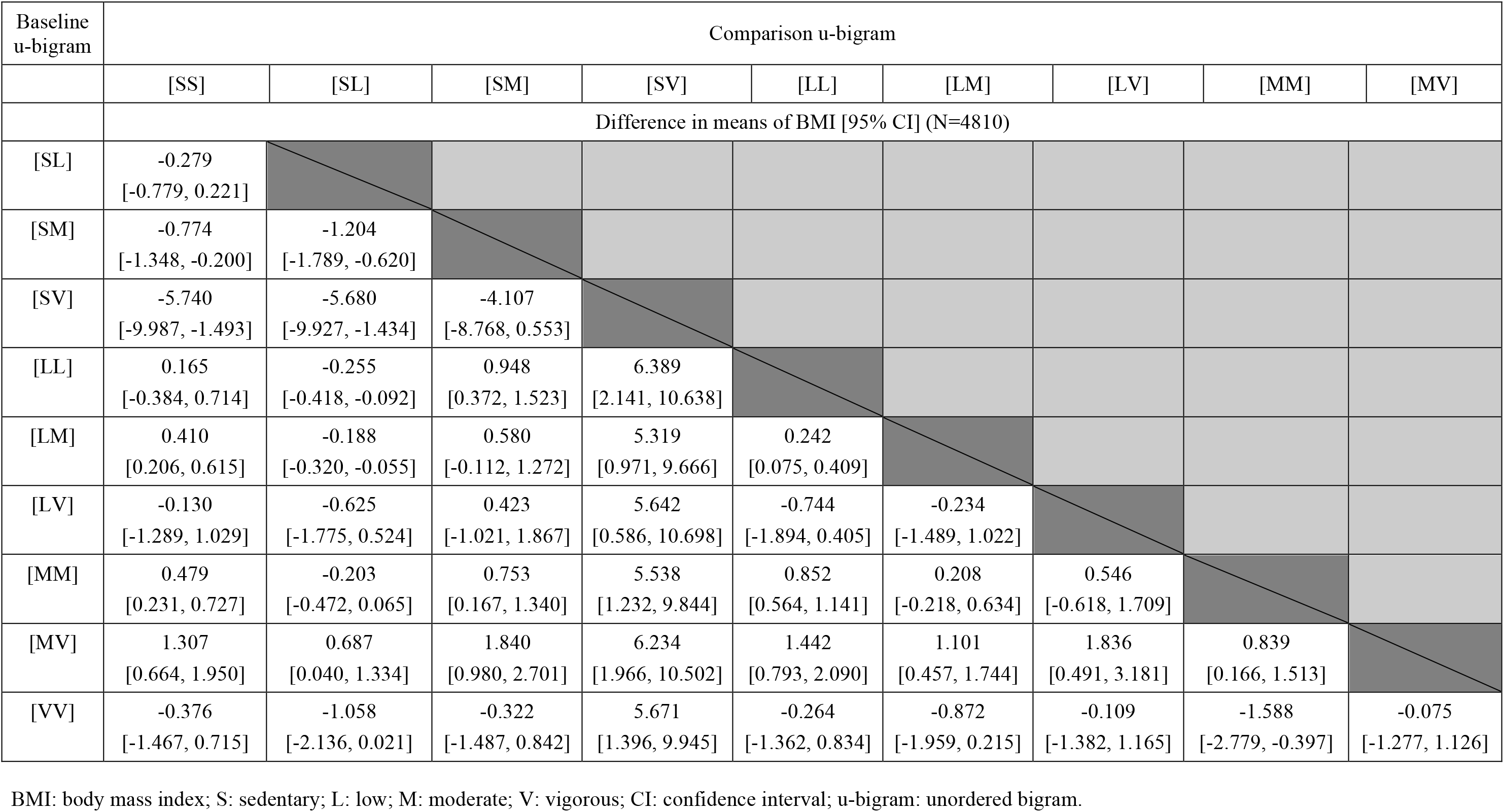
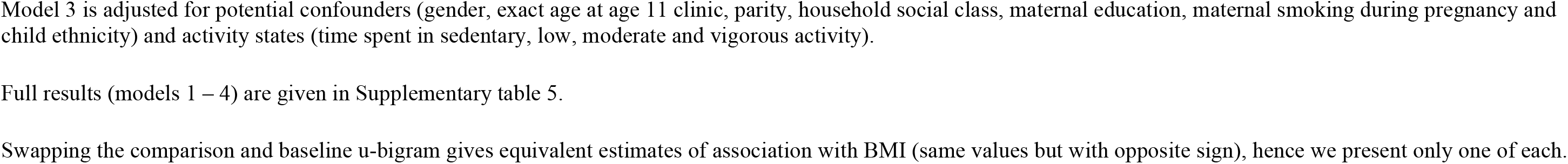
**Difference in means of BMI for a 10 epoch pair lower frequency of baseline u-bigram, coupled with a 10 epoch pair higher frequency of comparison u-bigram, after adjustment for confounding factors and amount of time spent in each activity state (model 3)**

| Baseline u-bigram | Comparison u-bigram |  |  |  |  |  |  |  |  |
| --- | --- | --- | --- | --- | --- | --- | --- | --- | --- |
|  | [SS] | [SL] | [SM] | [SV] | [LL] | [LM] | [LV] | [MM] | [MV] |
|  | Difference in means of BMI [95% CI] (N=4810) |  |  |  |  |  |  |  |  |
| [SL] | -0.279<br>[-0.779, 0.221] |  |  |  |  |  |  |  |  |
| [SM] | -0.774<br>[-1.348, -0.200] | -1.204<br>[-1.789, -0.620] |  |  |  |  |  |  |  |
| [SV] | -5.740<br>[-9.987, -1.493] | -5.680<br>[-9.927, -1.434] | -4.107<br>[-8.768, 0.553] |  |  |  |  |  |  |
| [LL] | 0.165<br>[-0.384, 0.714] | -0.255<br>[-0.418, -0.092] | 0.948<br>[0.372, 1.523] | 6.389<br>[2.141, 10.638] |  |  |  |  |  |
| [LM] | 0.410<br>[0.206, 0.615] | -0.188<br>[-0.320, -0.055] | 0.580<br>[-0.112, 1.272] | 5.319<br>[0.971, 9.666] | 0.242<br>[0.075, 0.409] |  |  |  |  |
| [LV] | -0.130<br>[-1.289, 1.029] | -0.625<br>[-1.775, 0.524] | 0.423<br>[-1.021, 1.867] | 5.642<br>[0.586, 10.698] | -0.744<br>[-1.894, 0.405] | -0.234<br>[-1.489, 1.022] |  |  |  |
| [MM] | 0.479<br>[0.231, 0.727] | -0.203<br>[-0.472, 0.065] | 0.753<br>[0.167, 1.340] | 5.538<br>[1.232, 9.844] | 0.852<br>[0.564, 1.141] | 0.208<br>[-0.218, 0.634] | 0.546<br>[-0.618, 1.709] |  |  |
| [MV] | 1.307<br>[0.664, 1.950] | 0.687<br>[0.040, 1.334] | 1.840<br>[0.980, 2.701] | 6.234<br>[1.966, 10.502] | 1.442<br>[0.793, 2.090] | 1.101<br>[0.457, 1.744] | 1.836<br>[0.491, 3.181] | 0.839<br>[0.166, 1.513] |  |
| [VV] | -0.376<br>[-1.467, 0.715] | -1.058<br>[-2.136, 0.021] | -0.322<br>[-1.487, 0.842] | 5.671<br>[1.396, 9.945] | -0.264<br>[-1.362, 0.834] | -0.872<br>[-1.959, 0.215] | -0.109<br>[-1.382, 1.165] | -1.588<br>[-2.779, -0.397] | -0.075<br>[-1.277, 1.126] |

BMI: body mass index; S: sedentary; L: low; M: moderate; V: vigorous; CI: confidence interval; u-bigram: unordered bigram.
Model 3 is adjusted for potential confounders (gender, exact age at age 11 clinic, parity, household social class, maternal education, maternal smoking during pregnancy and child ethnicity) and activity states (time spent in sedentary, low, moderate and vigorous activity).
Full results (models 1 – 4) are given in Supplementary table 5.
Swapping the comparison and baseline u-bigram gives equivalent estimates of association with BMI (same values but with opposite sign), hence we present only one of each.

While we present associations for a 10 frequency change, this may not represent feasible changes in activity for all u-bigrams, as their standard deviations vary widely (from 1.66 to 526.65 epoch pairs for the [SV] and [SS] u-bigrams, respectively; see Supplementary table 6). Hence, while the large estimates for [SV] in our main analysis appear unfeasible, this is because occurrences of [SV] are infrequent. Supplementary table 7 and Supplementary figure 18 presents associations for a 1 SD change of baseline bigram frequency, reflecting a realistic change based on variation in frequency of bigrams across our sample.

## DISCUSSION

In this work we have shown how activity bigrams can be used to investigate changes in activity from one moment to the next and how these can then be used to assess the associations of finely graded patterns of change in activity across a day with disease or health related traits such as BMI. Reciprocal bigrams (with the same sets of activity states, e.g. MV and VM) had comparable associations with BMI. This may be because these bigrams correlate highly with each other, as for example, people with more occurrences of the MV bigram on average have more VM bigrams.

Our tests of association of the u-bigrams with BMI identified several sequential activity patterns that were associated with BMI. In particular, a higher frequency of the [MV] u-bigram, coupled with a lower frequency of all other u-bigrams except [VV], was associated with a lower BMI, even after adjusting for mCPM and the amount of time spent in each activity state (sedentary, low, moderate and vigorous), respectively. This indicates that, given two groups of people who spent the same amount of time in each activity state per day and the same number of adjacent minutes in the vigorous state, those who have more adjacent minutes of moderate and vigorous, have a lower BMI. Hence, while current physical activity recommendations say adults should do at least 150 minutes of moderate-, or 75 minutes of vigorous-intensity physical activity a week (19,26,27), it may also be important to consider how activity levels change from one moment to the next. For instance, it might be that frequent occurrences of the acute increase in heart rate and changes to metabolism that occur with consecutive minutes in moderate and vigorous activity are important for lowering BMI.

Thus, if further research replicates our findings, demonstrates similar associations with other health related outcomes and evidence suggests these associations are causal then public health advice in relation to physical activity might need to change. To date, analysis and hence advice on physical activity has focused on average levels of activity. Exploring associations between sequential patterns of activity with other traits and disease will enable more comprehensive advice about the types and patterns of change in activity that may be beneficial or detrimental to health. For example, if a causal effect of adjacent minutes in moderate and vigorous activity on BMI was established and extended to obesity related disease outcomes such as diabetes and cardiovascular disease, it may be appropriate to advise performing moderate and vigorous activity sequentially, rather than separately throughout the day.

We also investigated whether the variance of activity levels (vCPM), was associated with BMI. While vCPM was associated with lower BMI, we found no evidence of an association independent of mCPM, indicating that the association of vCPM with BMI was due to its high correlation with mCPM.

### Study limitations

We use the average frequency of bigrams over 7 days as independent variables in linear regression models, without considering the uncertainty in these estimates. Therefore the confidence intervals of the associations with BMI may be under-estimated. Estimates based on a larger number of days may improve the accuracy of the bigram variables and hence the accuracy of associations based on these estimates. We identified differences in characteristics between ALSPAC participants included in our sample, and those who attended the age 11 clinic but were not included in our sample. This may bias associations if these data are not missing at random. However, the magnitudes of the differences were small and hence major bias unlikely. This paper is primarily concerned with demonstrating a novel (activity bigram) method. In future more applied papers we would want to undertake sensitivity analyses to explore the likelihood that our assumptions about missing data are correct.

We cannot infer that the associations with BMI we have presented in this paper are causal. Associations may be because the bigram (or u-bigram) has a causal effect on BMI. Alternatively, it may be the case that people with higher BMI are less likely to partake in activities that involve this type of activity pattern (i.e. more obese people may be less likely to change frequently from moderate to vigorous activity). Finally, while we adjusted for common confounding factors, it is possible that associations may be due to residual confounding.

We note that our analysis used a one-minute epoch such that the bigrams are a sequence of two one-minute intervals. The association of a bigram with another trait is likely to change as the epoch size changes. For example, a 1-minute epoch of moderate activity may be composed of one 30-second interval at low and one at vigorous activity, rather than continuous activity at the moderate level. While the accelerometers used in ALSPAC measured data in one-minute intervals, accelerometers are increasingly being used to collect raw data at a much higher resolution. Our methods are applicable to such data and could be used to determine whether associations with health/disease related traits differ with different epoch sizes. Also, while in this work we have used activity bigrams, it is possible to extend this approach to other n-grams. However, as *n* increases the number of occurrences of each n-gram in the population decreases and hence so does the study power.

To conclude, we have shown how a method initially developed for text data-mining can be used with accelerometer data to explore whether variation in physical activity intensity from one moment to the next, over and above mean levels of time spent in a given intensity, relates to health outcomes. We have shown that for BMI and activity bigrams calculated using a one-minute epoch, this does appear to be the case. We recommend that other studies explore whether our findings with BMI replicate, and the association of activity bigrams with other traits are assessed.

## Acknowledgements

We are extremely grateful to all the families who took part in this study, the midwives for their help in recruiting them, and the whole ALSPAC team, which includes interviewers, computer and laboratory technicians, clerical workers, research scientists, volunteers, managers, receptionists and nurses. We are grateful to the GW4 Consortium: “Analysis of Intensively-Collated Health Data” workshops for providing a platform for discussion and knowledge sharing (funded by GW4 grant GW4-IF4-008).

